# Target-dependent siRNA suppression distinguishes self from non-self endogenous siRNAs in *C. elegans* germline

**DOI:** 10.1101/2022.01.25.477739

**Authors:** Zoran Gajic, Diljeet Kaur, Julie Ni, Zhaorong Zhu, Anna Zhebrun, Maria Gajic, Matthew Kim, Julia Hong, Monika Priyadarshini, Christian Frøkjær-Jensen, Sam Gu

**Author notes:** These authors contributed equally to this work (Zoran Gajic, Diljeet Kaur, Julie Ni).

## Abstract

Despite their prominent role in transposon silencing, expression of endo-siRNAs is not limited to the “non-self” DNA elements. Transcripts of protein-coding genes (“self” DNA) in some cases also produce endo-siRNAs in yeast, plants, and animals [1]. How cells distinguish these two populations of siRNAs to prevent unwanted silencing of self-genes in animals is not well understood. To address this question, we examined the expression of ectopic siRNAs from an LTR retrotransposon in *C. elegans* germline. We found that the abundance of ectopic siRNAs was dependent on their homologous target genes: ectopic siRNAs against genes expressed only in somatic cells can be abundantly expressed. In contrast, ectopic siRNAs against germline-expressed genes are often suppressed. This phenomenon, which we termed “target-directed siRNA suppression”, is dependent on the target mRNA and requires germline P-granule components. We found that siRNA suppression can also occur to naturally produced endo-siRNAs. We suggest that siRNA suppression plays an important role in regulating siRNA expression and preventing self-genes from aberrant epigenetic silencing.

## Introduction

Small RNAs, such as microRNAs, endo-siRNAs, and piRNAs carry out a diverse set of cellular functions through their ability to silence homologous genes. Small RNAs orchestrate gene silencing by serving as guide molecules to bring Argonaute and other regulatory proteins to the target RNA transcripts. Therefore, the steady state level of small RNAs is a key determinant of the gene silencing activity.

The endo-siRNA pathway in *C. elegans* germline is a powerful system to explore small RNA biology and transgenerational epigenetic inheritance [2]. Among the different classes of endo-siRNAs, the ones that target transposons and other repetitive DNA elements belong to secondary siRNA or 22G-RNA class [3], which are *de-novo* synthesized by RNA-dependent RNA polymerases (RdRPs) using the target mRNAs as the template. 22G-RNA synthesis can be triggered by a diverse set of RNA molecules/structures: dsRNA [4], 26G-RNA [5, 6], piRNA [7, 8], aberrant mRNA processing [9, 10], and untemplated RNA-tailing [11]. Once produced, these siRNAs are bound by the germline AGO proteins such as WAGO-1 [3] and HRDE-1 [12], which induce post-transcriptional and transcriptional repression at the target genes. The steady state level of siRNAs is determined through both biogenesis and turnover of siRNAs. Recent studies have identified numerous biochemical activities that can affect the siRNA turnover rate, such as siRNA tailing [13-16] and the stability and processing of AGO proteins [17, 18]. In these cases, it is unclear to what extent the siRNA turnover is mediated by their target mRNA.

Interestingly, actively transcribed germline genes also naturally produce 22G-class siRNAs in *C. elegans* [19, 20]. For the purpose of this study, we refer to these siRNAs as self siRNAs and the ones from transposons as non-self siRNAs. Both populations are synthesized by RdRPs and share the same size (22 nt) and 5’ guanine biases. Despite these similarities, the self siRNAs differ from the non-self siRNAs in at least two aspects. Firstly, self siRNAs have much lower density, as measured by normalized read count per unit length of mRNA, than non-self siRNAs (Fig. S1). Secondly, they are enriched in different AGO proteins: the non-self siRNAs in WAGO-1 and HRDE-1 and self siRNAs in another germline-specific AGO protein CSR-1 [19]. Self siRNAs have been suggested to fine-tune germline gene expression: loss of the germline RdRP enzyme EGO-1 leads to reduced siRNA and increased mRNA expressions of germline genes [20]; loss of the CSR-1 protein also led to complex dysregulation of germline gene expression and ultimately sterility [19, 21-25].

Despite the clear distinctions in their abundance and biochemical properties, how cells distinguish the self and non-self siRNAs remains largely unknown. Transposons and self genes differ significantly in their chromatin environment, modes of transcription, RNA processing and trafficking, which can all potentially affect siRNA production and loading into AGOs. However, it is hard to change one factor without affecting other. In this study, we took a genome engineering approach to express ectopic siRNAs from a natural “hot spot” of non-self siRNAs. This enables us to examine the fate of the non-self and ectopic self siRNAs expressed from the same genomic locus.

## Results

### The design of ectopic siRNA production from the LTR retrotransposon *Cer3*

In this study we used CRISPR to insert approximately 400 nt exonic sequences from various protein-coding genes into the LTR retrotransposon *Cer3* to test its ability to express ectopic siRNAs (Fig. 1A). *Cer3* is a native target of germline nuclear RNAi and produces abundant germline-specific siRNAs (Fig. 1B and Fig. S2) [26, 27]. There is only one copy of *Cer3* in the genome of the wild type Bristol N2 strain. The insertion site was chosen for its local peak level of siRNA production (Fig. S2). Single nucleotide polymorphisms (SNPs) were introduced in some of the insertions at 30 nt intervals to distinguish between the transposon-driven ectopic siRNAs and the native self siRNAs (generated from the homologous target genes).

**Figure 1.**
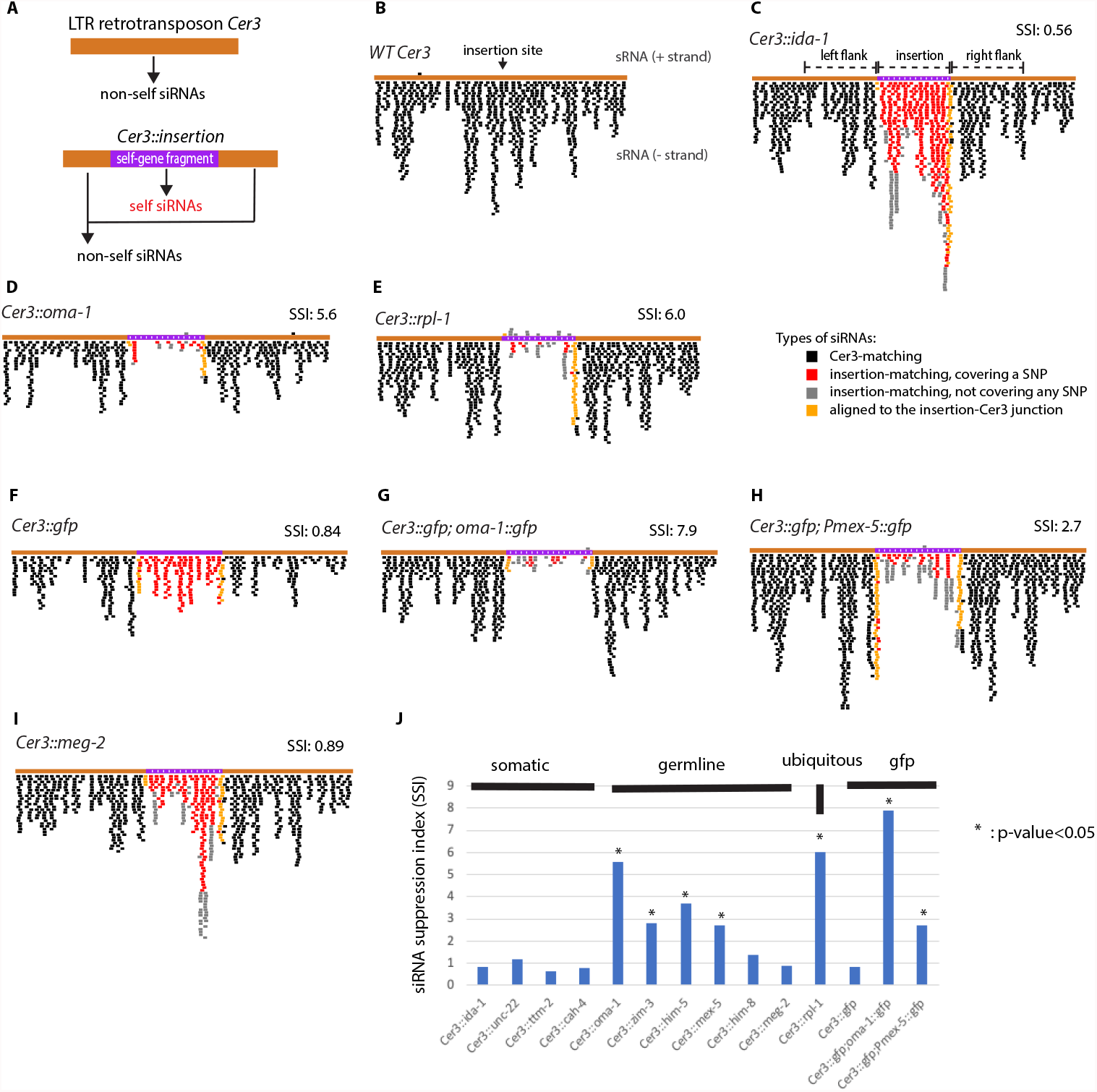
Differential expression of ectopic self siRNAs driven by LTR retrotransposon *Cer3*. (**A**) A schematic diagram of *Cer3* with a CRISPR-engineered insertion (*Cer3::insertion*) to express ectopic siRNAs. (**B**-**I**) siRNA track plots for WT *Cer3* and various *Cer3::insertion* alleles. Only the insertion and the 1.4 kb *Cer3* sequence (700 bp on either side of the insertion) are included in the plots. Sense and antisense small RNAs with perfect alignment to the *Cer3::insertion* sequences are plotted above and below the gene track, respectively. siRNA tracks are color-coded to reflect their locations as indicated in the legend. Additional *Cer3::insertion* siRNA track plots are in Fig. S4. (**J**) Bar graph of siRNA suppression indexes for various *Cer3* alleles shown in Fig. 1 and S4. The siRNA suppression index (SSI=*Cer3* siRNA density of the 400 bp left and right flanking sequences / siRNA density of the insertion) and p-value (Wilcoxon Rank Sum Test, null hypothesis: the density of siRNAs mapped to the insertion is the same or larger than the density of flanking *Cer3* siRNAs) are also indicated for each panel.

### Suppression of *Cer3*-driven ectopic siRNAs against germline genes

Our sRNA-seq analysis indicated that none of the insertions used in this study affected the *Cer3* siRNA expression (Fig. 1 and S3). However, the levels of the ectopic siRNAs from different insertions varied significantly. Insertions of the *gfp* fragment (*Cer3::gfp*) (Fig. 1F) and four somatic gene fragments (*ida-1, ttm-2, cah-4, and unc-22*, selected for preferentially neuronal, intestinal, hypodermal, or muscle expression, respectively based on RNA-seq data from [28]) (Fig. 1C, Fig. S4A-C) produced abundant siRNAs, with levels similar to the ones expressed from the flanking *Cer3* sequences (Fig. 1J). In contrast, fragments from four germline-expressed genes (*oma-1, zim-3, him-5*, and *mex-5*) and one ubiquitously expressed ribosomal protein gene (*rpl-1*) produced significantly fewer siRNAs than the flanking *Cer3* siRNAs (Figs. 1D-E, S4D, S4G, 1J). We also crossed the *Cer3:gfp* allele into strains that carry a germline-expressed *gfp* transgene, either a translational fusion of *oma-1::gfp* or *gfp* driven by a germline-specific promoter (*Pmex-5::oma-1*). We found that the *Cer3*-driven *gfp* siRNAs were produced in much lower abundance in both the *gfp* transgene (+) animals than in the *gfp* transgene (-) animals (Fig. 1G and H). Of the nine germline-expressed fragments inserted into *Cer3*, we note that two (*meg-2* and *him-8*) were not associated with repressed ectopic siRNA expression (Fig. 1I and S4F). We do not know the reason for these exceptions at this point (see more in Discussion).

To compare the relative expression of siRNAs from *Cer3* and insertions, we defined an siRNA suppression index (SSI) as the ratio between the density of flanking *Cer3* siRNAs and insertion siRNAs. The average SSI for the germline-expressed fragments (except *him-8* and *meg-2*) fragments was 4.5. In contrast, the average SSI for *gfp* (WT background) and somatic fragments is 0.8. Suppression of ectopic siRNAs also occurred when we inserted the same *oma-1* fragment into a second LTR retrotransposon *Cer8* (Fig. S4 H and I). These results indicate that transposon-driven self siRNAs with homology to germline-expressed genes tend to be suppressed. We refer to this phenomenon as “siRNA suppression” in this paper.

### siRNA suppression requires the homologous target gene

The germline gene *oma-1* has been extensively used as a native gene to study RNAi and transgenerational epigenetic silencing [29]. For the rest of the study, we used the *Cer3::oma-1* allele to characterize siRNA suppression. To determine whether the siRNA suppression at *Cer3::oma-1* requires the endogenous *oma-1* DNA sequence, we crossed the *Cer3::oma-1* allele into two *oma-1* mutants: (1) a

1.5 kb deletion (*oma-1[tm1396]*) [30] including the promoter and a large fraction of the transcribed sequence of *oma-1*, and (2) a smaller deletion (*oma-1[red110]*), generated for this study, which deletes a 240 bp fragment within the 400 bp homologous sequence to the *Cer3::oma-1* insertion. We found that *oma-1(tm1396)* abrogated the siRNA suppression effect for the entire *oma-1* insertion of *Cer3::oma-1* (Fig. 2A). *oma-1(red110)* abrogated siRNA suppression only for the sequence that is homologous to the deletion (Fig. 2B). These results, together with the ones mentioned in the previous section, indicate that (1) the ectopic siRNA suppression requires the homologous DNA sequence in the target gene and (2) the suppression is highly local and does not spread to flanking non-homologous sequence in *Cer3*.

**Figure 2.**
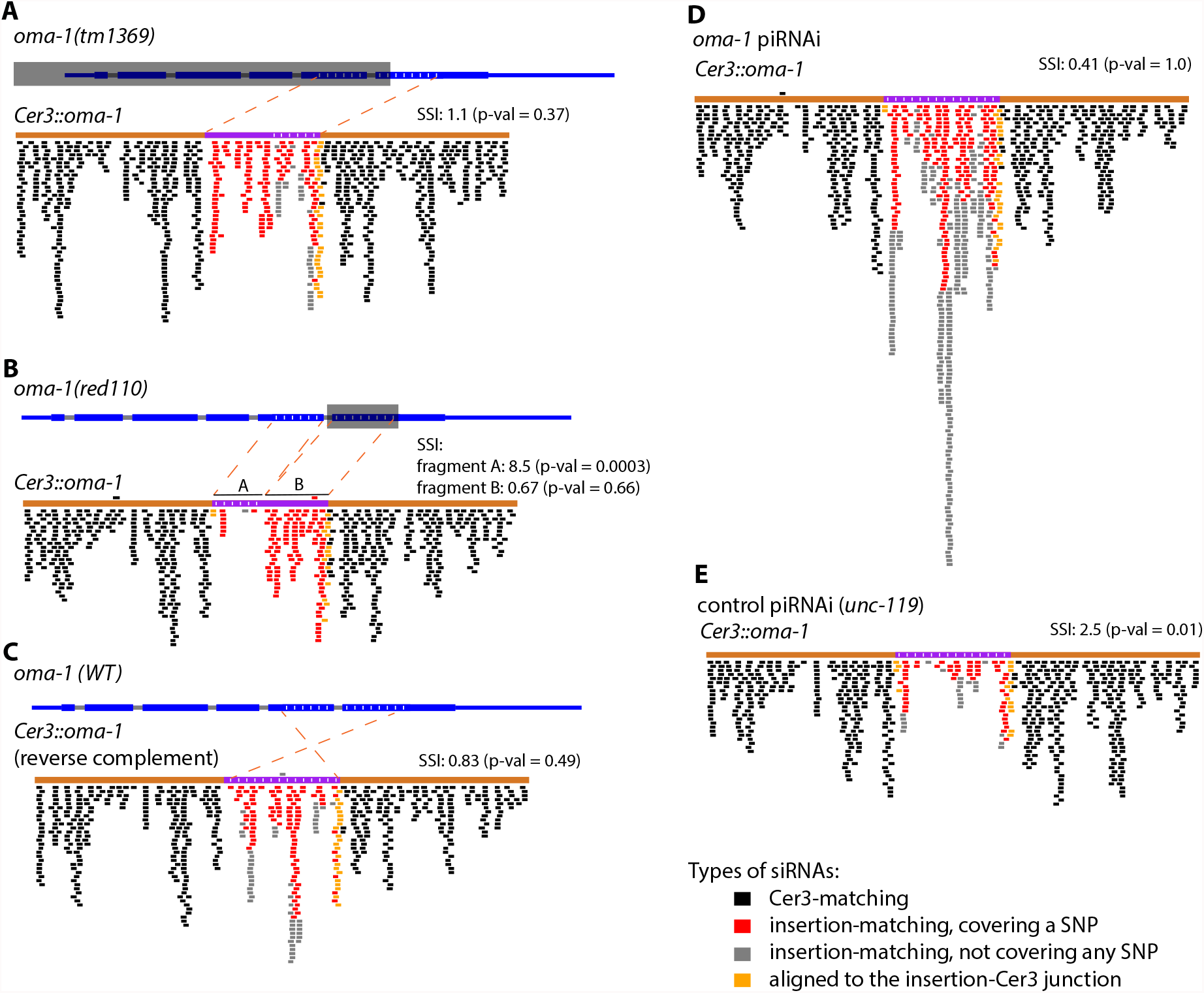
siRNA suppression requires the mRNA expression of the homologous gene. (**A**-**B**) *Cer3::oma-1* siRNA track plots for strains carrying *oma-1* deletion mutations (A: *tm1396*, B: *red110*). Deleted regions are indicted by transparent grey boxes. (**C**) A siRNA track plot for *Cer3::oma-1* that expresses sense-stranded *oma-1* siRNAs. (**D**-**E**) siRNA track plots for *Cer3::oma-1* in *oma-1* piRNAi animals and in control animals (*unc-119* piRNAi). See Fig. S5 for piRNAi-induced *oma-1* mRNA silencing and Fig. S6 for additional experiments of *oma-1* piRNAi.

### siRNA suppression is likely mediated by mRNA

We hypothesized that siRNA suppression is mediated by the target mRNA and thus tested the strand specificity and the effect of mRNA silencing on siRNA suppression. For strand specificity, we compared two *Cer3::oma-1* alleles that differ in the orientation of insertion so that one produces antisense siRNAs against *oma-1* mRNA (Fig. 1D) and the other sense siRNAs against *oma-1* mRNA (Fig. 2C). We found that, in contrast to antisense ectopic siRNAs, sense-stranded ectopic siRNAs were not suppressed (Fig. 2C). Therefore, the siRNA suppression effect is specific to antisense siRNAs.

To test the effect of silencing *oma-1* on siRNA suppression, we first performed *oma-1* RNAi by dsRNA-feeding and found that the siRNA suppression was still observed for *Cer3::oma-1* (Fig. S5A). We then performed a piRNA-triggered *oma-1* silencing (piRNAi) using a piRNA-expressing transgene approach recently developed in [31]. Consistent with the previous report [31], our RNA-seq analysis indicated that piRNAi induced a much more robust *oma-1* silencing (29.1-fold) than dsRNA (4.6-fold) (Fig. S5B and C). *oma-1* piRNAi abrogated the siRNA suppression for *Cer3::oma-1* (Fig. 2D and S6A). Together, these results suggest that siRNA suppression is mediated by the mRNA of the homologous germline gene.

### siRNA suppression does not require the germline AGO proteins HRDE-1, CSR-1, or the piRNA pathway, but requires the P-granule components

We then performed a small scale candidate gene-based screen to investigate the genetic requirement of siRNA suppression using *Cer3::oma-1*. CSR-1 and HRDE-1 are two germline-specific AGO proteins that preferentially bind different populations of germline siRNAs: self siRNAs for CSR-1 and non-self siRNAs for HRDE-1 [7, 12, 19, 32]. We examined siRNA expressions of *Cer3::oma-1* in the *hrde-1(-)* animals and CSR-1-depleted animals by auxin-induced protein degradation, and found that neither HRDE-1 or CSR-1 was required for siRNA suppression (Fig. 3B-C, S7).

**Figure 3.**
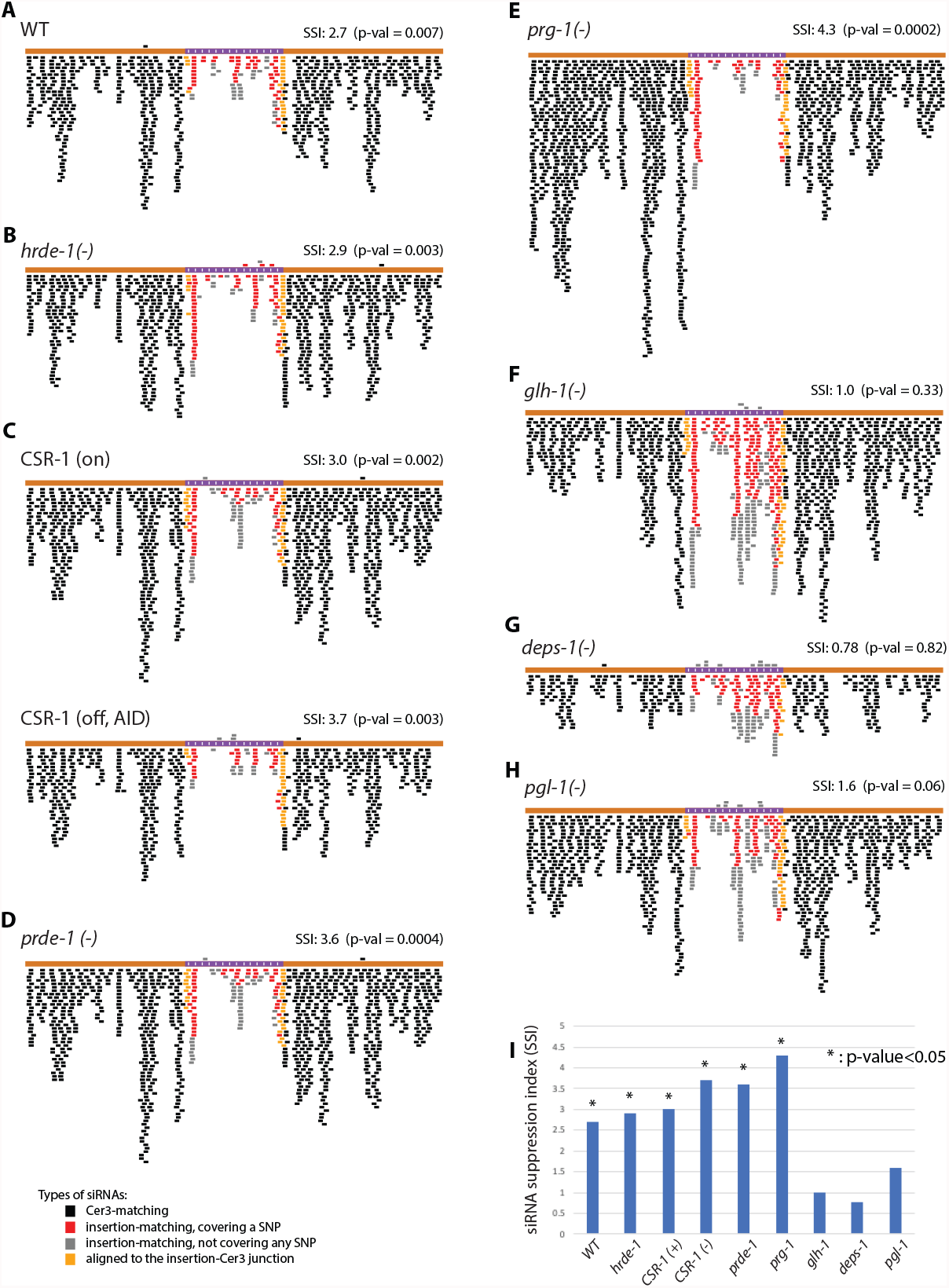
Genetic requirement of siRNA suppression of *Cer3::oma-1*. (**A-B, D**-**H**): *Cer3::oma-1* siRNA track plots for wild type animals and various loss-of-function mutants as indicated in each panel. (**C**) *Cer3::oma-1* siRNA track plots for animals with auxin-induced degradation (AID) of CSR-1 (lower panel) and control animals (upper panel). See Fig. S7 for CSR-1 AID. (**I**) Bar graph of siRNA suppression indexes (SSI) shown in panels A-H. See Fig. 1 legend for the SSI and p-value calculations.

Recent studies have shown that the piRNA pathway can suppress the run-away siRNA amplification in *C. elegans* [11, 33, 34]. The PRDE-1 protein is required for the biogenesis of piRNAs [35, 36] and PRG-1 is the PIWI protein that binds piRNAs [18]; both are essential for the piRNA-mediated functions [2]. We observed strong siRNA suppression of *Cer3::oma-1* in the *prg-1(-)* and *prde-1(-)* animals (Fig. 3D and E), indicating that piRNA activity is not required for siRNA suppression of *Cer3::oma-1*.

The P granules in *C. elegans* germline are liquid-like, membrane-less condensates of RNA and proteins, that often locate adjacent to the cytoplasmic side of the nuclear pore complexes [37]. Many proteins in the RNAi pathway are enriched in the P granules, and the P granules have been shown to promote RNAi [38, 39]. Consistent with this notion, we found that mutant animals lacking any of the P-granule assembly factors DEPS-1, GLH-1, and PGL-1 showed reduced levels of *Cer3* siRNAs (Fig. S8). We crossed the *Cer3::oma-1* allele into these P-granule mutants, and found that the siRNA suppression effect was abrogated in *deps-1(-)* and *glh-1(-)* animals (Fig. 3F-G). The *pgl-1(-)* mutation also reduced the siRNA suppression effect albeit to a lesser degree than *deps-1(-)* or *glh-1(-)* (Fig. 3H). These results suggest that siRNA suppression requires functional P granules.

### Unsuppressed ectopic siRNAs induce transitive RNAi of the target gene

To determine the impact of *Cer3*-driven ectopic siRNAs on the target gene mRNA expression, we performed RT-qPCR analyses of the corresponding target genes in worms carrying the *Cer3::oma-1, Cer3::zim-3, Cer3::rpl-1* or *Cer3::meg-2* allele. For the *Cer3::oma-1, Cer3::zim-3*, and *Cer3::rpl-1* alleles, which all exhibited strong siRNA suppression, their corresponding target gene mRNAs were expressed at the wild-type level (Fig. 4 A-C), and their siRNA levels were not affected either (Fig. 4E-G), indicating a lack of RNAi at these genes. In contrast, *Cer3::meg-2*, which is resistant to siRNA suppression, was associated with a 43% reduction in *meg-2* mRNA and a 28-fold increase in *meg-2* siRNAs. These results indicate that the *Cer3*-driven ectopic siRNAs, if not suppressed, can induce a transitive RNAi effect at the target gene.

**Figure 4.**
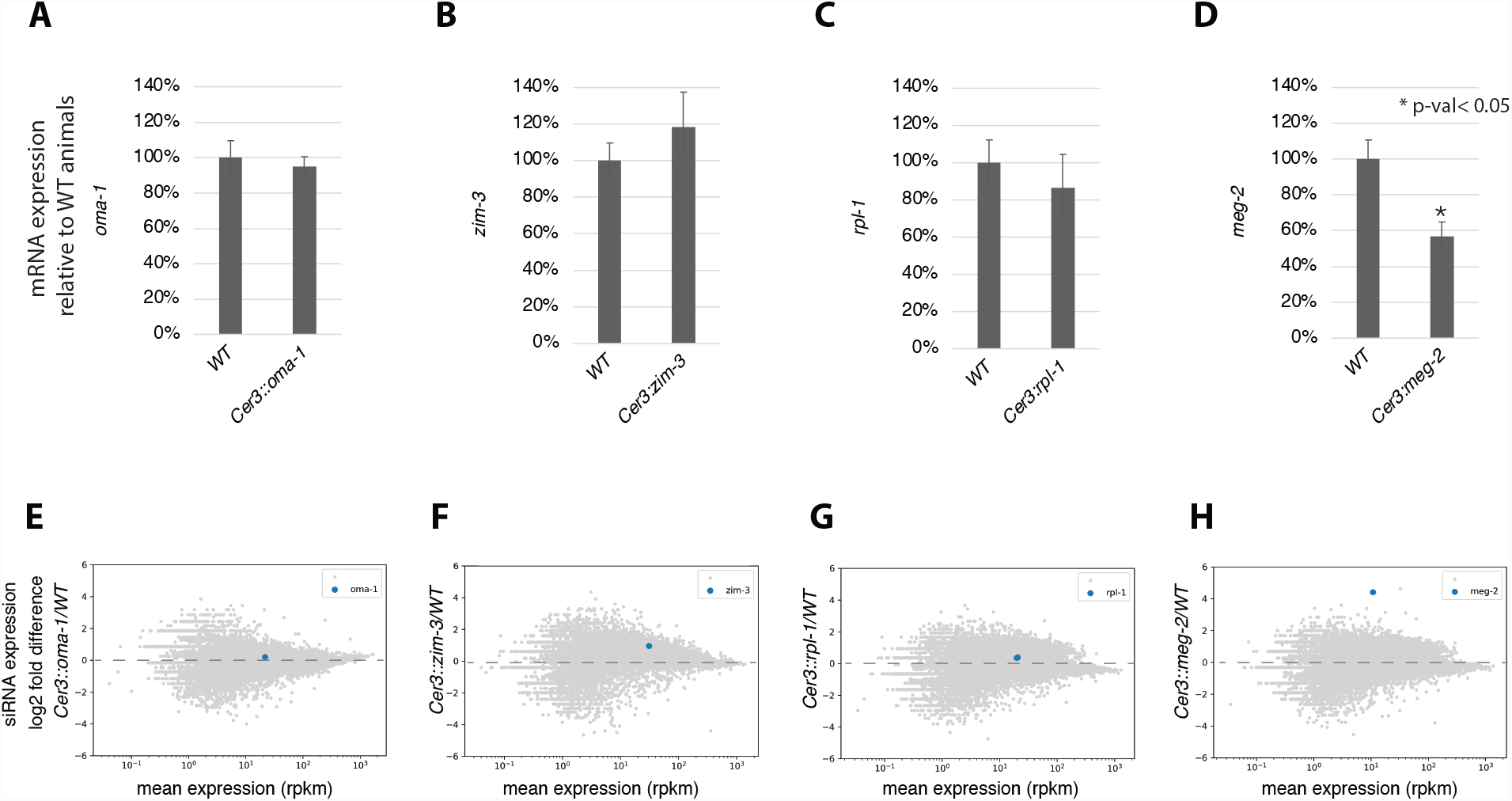
The impact of *Cer3-*driven ectopic siRNAs on target gene mRNA and siRNA expressions. (**A**-**D**) RT-qPCR analysis comparing WT and *Cer3* mutant animals for the corresponding target gene mRNA expression. (**E**-**H**) sRNA-seq analysis (MA-plots) comparing siRNA expression in WT and *Cer3* mutant animals. The corresponding target gene in each panel is highlighted.

### siRNA suppression of the native siRNAs

So far, our experiments only examined ectopic siRNAs expressed from genetically engineered loci. We next wished to determine whether the native siRNAs are sensitive to siRNA suppression. Native germline nuclear RNAi targets have the hallmarks of low mRNA and high siRNA expressions. We took two different approaches to test whether native non-self siRNAs can be suppressed by a high level of homologous mRNAs.

First, we inserted a *Cer3* fragment in the 3’ UTR of the native *oma-1* gene (*oma-1::Cer3*) (Fig. 5A). The *Cer3* fragment was chosen for its high siRNA expression in *Cer3*. The *Cer3* insertion did not significantly affect the expressions of *oma-1* mRNA or siRNA (Fig. S9A-B). In the *oma-1::Cer3* animals, the expression of the homologous siRNAs from *Cer3* was drastically suppressed, and the effect was specific to the homologous sequence without spreading to the adjacent *Cer3* sequence (Fig. 5A-B and Fig. S9C).

**Figure 5.**
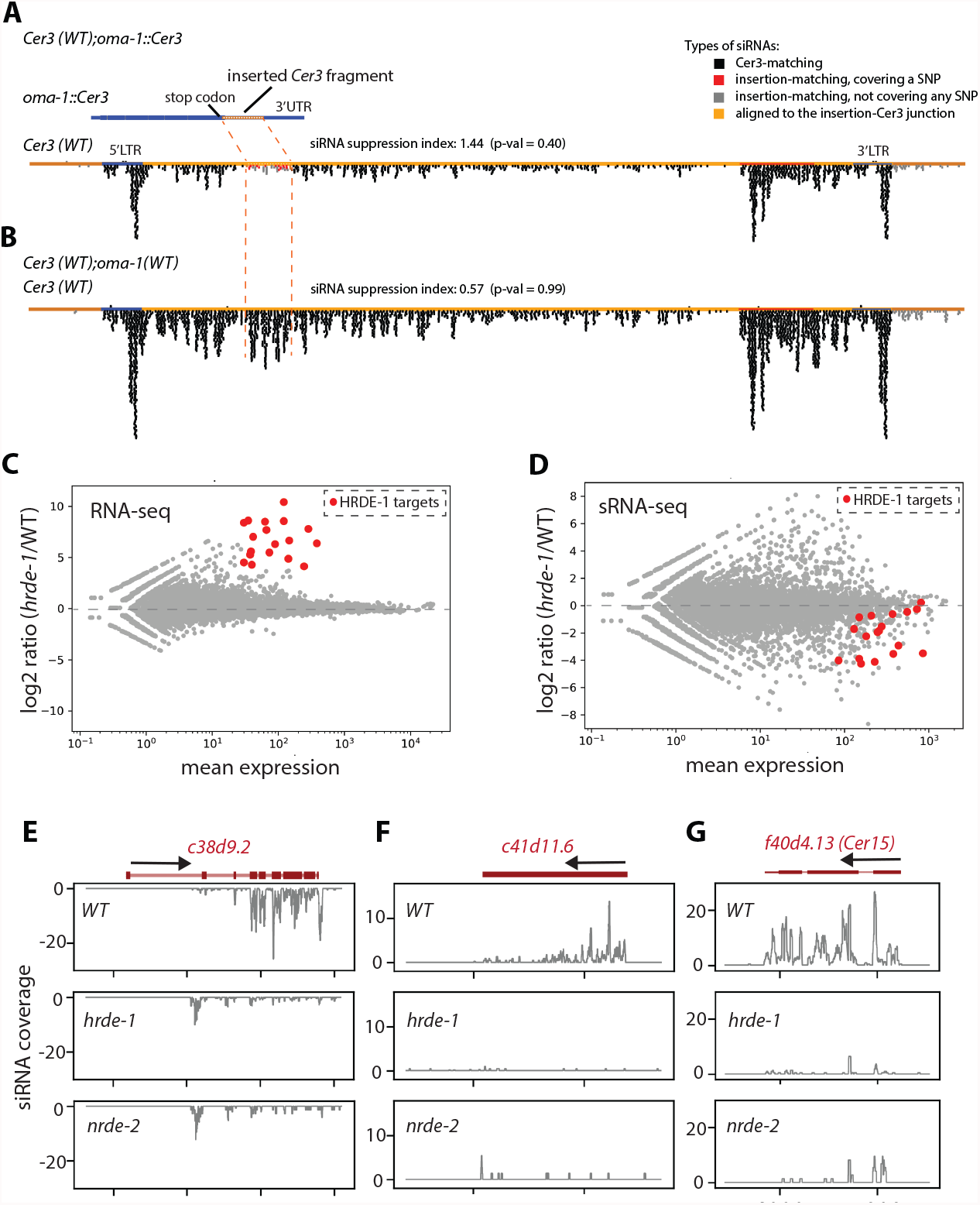
Suppression of native non-self siRNAs. (**A**) siRNA track plot of a full length WT *Cer3* in the *oma-1::Cer3* mutant, in which a 467 nt *Cer3* sequence was inserted immediately after the stop codon of *oma-1*, with SNPs every 30 nt. (**B**) *Cer3* siRNA track plot for WT animals. (**C**-**D**) MA(log2 ratio versus mean average)-plots comparing *hrde-1* and WT RNA-seq (C) and sRNA-seq (D). Top 20 HRDE-1 target genes were highlighted in red. (**E**-**G**) sRNA coverage plots for three different HRDE-1 targets in WT, *hrde-1*, and *nrde-2* mutants, as indicated.

In our second experiment, we asked whether the desilencing of native HRDE-1 targets can lead to their own siRNA suppression. To this end, we analyzed our previously published mRNA-seq and sRNA-seq data of the WT and *hrde-1*(-) animals cultured at 23°C. We chose this temperature because *hrde-1(-)* animals have enhanced desilencing at a high temperature [40]. Out of the top 20 desilenced genes in the *hrde-1(-)* mutant (>16-fold desilencing, p<0.05), 12 genes had at least 3-fold decreases in siRNA expression (p<0.05). Similar siRNA losses were observed in another nuclear RNAi-defective mutant *nrde-2* [41] (Fig. 5E-G and data not shown).

These results indicate that siRNA suppression is not limited to genetically engineered ectopic siRNAs, but can also occur to native siRNAs when the homologous mRNA sequence is actively expressed *in cis* or *in trans*.

## Discussion

In this study we developed a strategy to express ectopic siRNAs from a native germline endo-siRNA hotspot (*Cer3*) in the *C. elegans* genome. In the absence of any germline-expressed target gene, the ectopic siRNAs can be abundantly expressed. In the presence of germline-expressed target gene, we found that the steady state level of ectopic siRNAs can be suppressed by homologous mRNAs. Previous studies have shown that factors that promote siRNA turnover play a key role in preventing unwanted silencing in the genome [13-16]. In these cases, it was unclear to what extent the siRNA turnover was dependent on the target RNA. Our study demonstrates that target-direct siRNA suppression is an integral component of the RNAi pathway in *C. elegans* germline and plays an important role in distinguishing self and non-self genetic material.

### Potential biological functions

RNAi in *C. elegans* germline is highly robust and long-lasting [4, 29, 42-44]. These features, while essential for genome surveillance against non-self DNA, can potentially lead to unwanted silencing of self genes. Previous studies have shown that epimutations of self genes can be induced by a diverse set of experimental triggers and genetic conditions [7, 11, 33, 34, 45-47]. The risk of epimutation is further increased by the presence of the siRNAs that are naturally produced from germline genes. For example, rRNA genes appear to be particularly prone to aberrant siRNA production and silencing [15, 33, 34]. Some active genes contain siRNA-producing transposon elements in their introns or nearby intergenic regions. In addition, a recent epimutation accumulation study found that siRNAs can increase for certain self-genes in wild type populations [48]. Interestingly, such increases appear to be transient. These observations highlight the importance of regulating self-targeting siRNAs.

The aberrant RNAi of self genes may be prevented by multiple mechanisms. The lack of silencing triggers, such as dsRNA, piRNA, 26G-RNA, pUG or other untemplated tails, is likely a major reason for the absence or low level of siRNAs from self genes [2]. siRNA suppression also occurs in *S. pombe* to control the unwanted RNAi [14]. The target-dependent siRNA suppression can provide another mechanism of distinguishing the self and non-self siRNAs. Given the large difference in the mRNA levels between the self and non-self genes, the dependance on the target mRNA can ensure the specificity of self-siRNA suppression and avoid suppressing the non-self siRNAs. Target-directed siRNA suppression may also contribute to the previously observed non-coding function of mRNA in promoting gene expression [22, 44, 46, 49]. We note that target-directed siRNA suppression does not completely abolish ectopic siRNAs. Rather this feature is perhaps important for the fine tuning the siRNA levels of germline genes.

### Mechanistic considerations

#### siRNA synthesis or degradation?

In principle, there are two ways to achieve target-directed siRNA suppression: inhibiting siRNA biogenesis or enhancing siRNA turnover. At this point we find it difficult to imagine how target mRNAs *directly* inhibit the synthesis of ectopic siRNA synthesis. siRNAs produced from the target mRNA, on the other hand, can potentially bind to the homologous RNA sequence inserted in *Cer3*, and function as a barrier against the RdRP activity, as suggested by [50]. However, such model would predict that the siRNAs from the upstream *Cer3* sequence that flanks the insertion would be suppressed as well. We did not observe such effect. Instead, the siRNA suppression was specifically limited to the homologous sequence. Although we cannot rule out a model involving siRNA synthesis inhibition, we currently favor a target mRNA-mediated siRNA degradation model, perhaps using a mechanism that is similar to target-dependent microRNA degradation [51, 52] or RNA tailing mediated sRNA degradation [13-16, 53]. In addition, the steady state level of siRNAs can also be affected by activities that influence the Argonaute proteins’ ability to bind siRNAs [17]. We did not observe any above-background level of tailing for *Cer3*-driven ectopic siRNAs (data not shown), but we cannot rule out the possibility of rapid siRNA degradation after tailing. Recent studies have shown that the piRNA pathway, in addition to its silencing role, protect the rRNA locus, histones, and other self genes from aberrant siRNA production and silencing [11, 31, 33, 34]. Future studies are needed to determine to what extend these activities are directed by target RNA.

#### P granule

We found that loss of P-granule components GLH-1, DEPS-1, or PGL-1 leads to siRNA suppression defect. Future studies are needed to test whether siRNA suppression occurs in the P granules. Such possibility is intriguing in that the P granules and other adjacent paranuclear condensates have been suggested as a hub for siRNA production and mRNA aggregation [39]. The close proximity of siRNA biogenesis and siRNA suppression could reduce the chance of the unwanted siRNAs escaping from the quality control mechanism. One complication is that mutations in *glh-1, deps-1*, and *pgl-1* also reduce endo-siRNA production at *Cer3* and elsewhere, which compromises the utility of these mutants in studying the function of siRNA suppression.

#### Complex relationship between mRNA and siRNA

One paradox of RNAi is that the transcripts are both the target and a necessary component of RNAi by being the templates of siRNA biogenesis. Our study shows that the target transcripts can also suppress the homologous siRNAs, which further increases the complexity to the mRNA-siRNA relationship. Future studies are needed to study how such seemingly opposing activities are coordinated/separated either chemically, spatially, or temporally.

#### The exceptions

Our study showed that the germline expression of the target mRNA is a necessary factor but not sufficient determinant in siRNA suppression. Different modes of transcriptional and post-transcriptional regulation, sub-cellular localization of mRNAs, and other factors may influence the utility of germline gene in target-directed siRNA suppression. Future studies are needed to identify additional rules of siRNA suppression.

We suggest that the target-directed siRNA suppression may distinguish self and non-self siRNA and play additional functions in other eukaryotes. We also note that such activity should be considered in mRNA-based technology and therapy.

## Supplementary figure legend

**Fig. S1:**
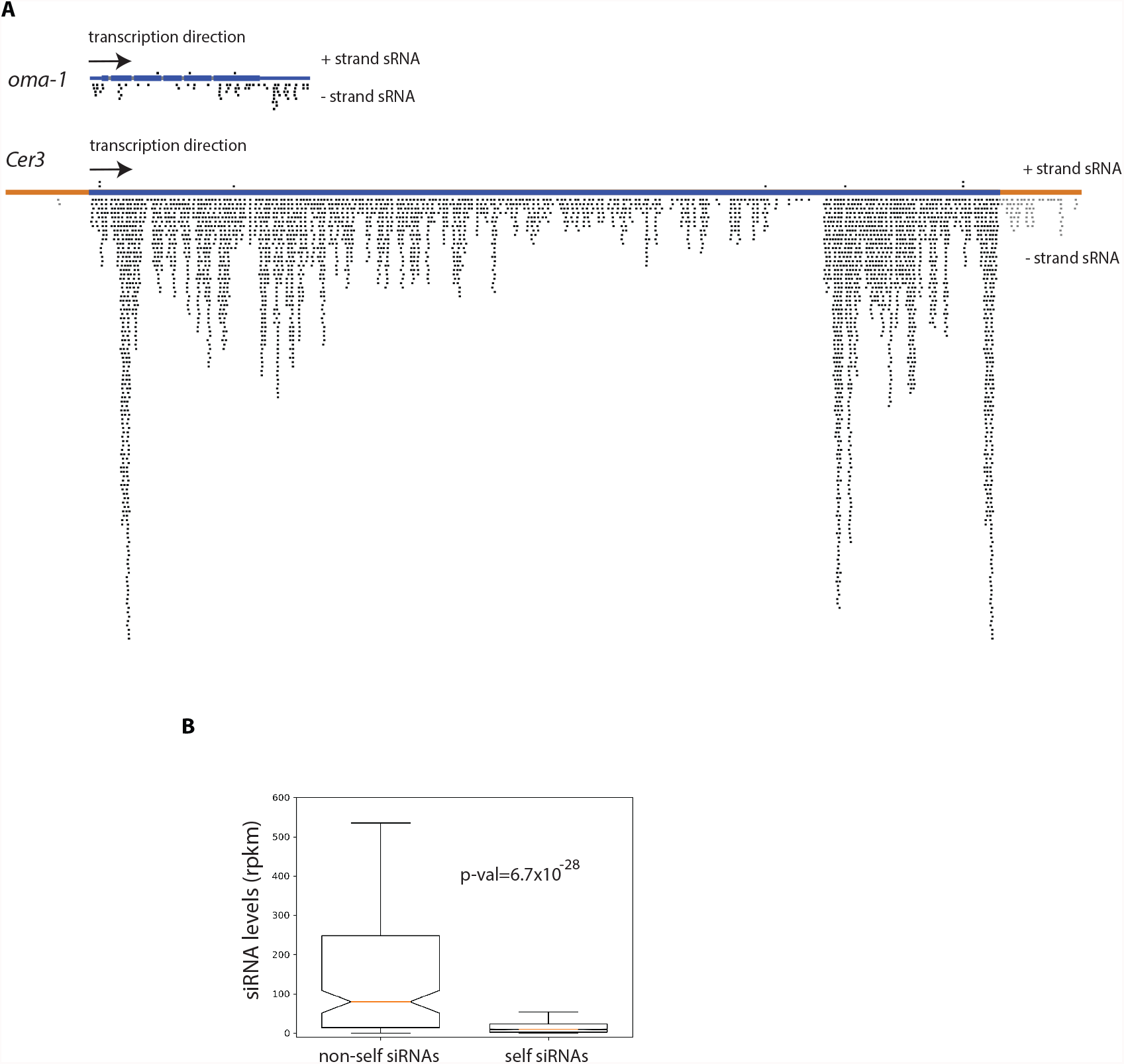
Differential expression of self and non-self siRNAs. (**A**) siRNA track plots for *oma-1* and the LTR retrotransposon *Cer3* in WT adult animals from the same sequencing run. Each sequenced small RNA read is indicated as a black block above (sense sRNA) or under (antisense sRNA) the gene track. (**B**) Box plot of average siRNA levels of native germline nuclear RNAi targets (non-self siRNAs) and germline genes (self siRNAs) in WT animals. The native germline nuclear RNAi target genes were obtained from [26]. The germline genes are the oogenic genes identified in [54].

**Fig. S2:**
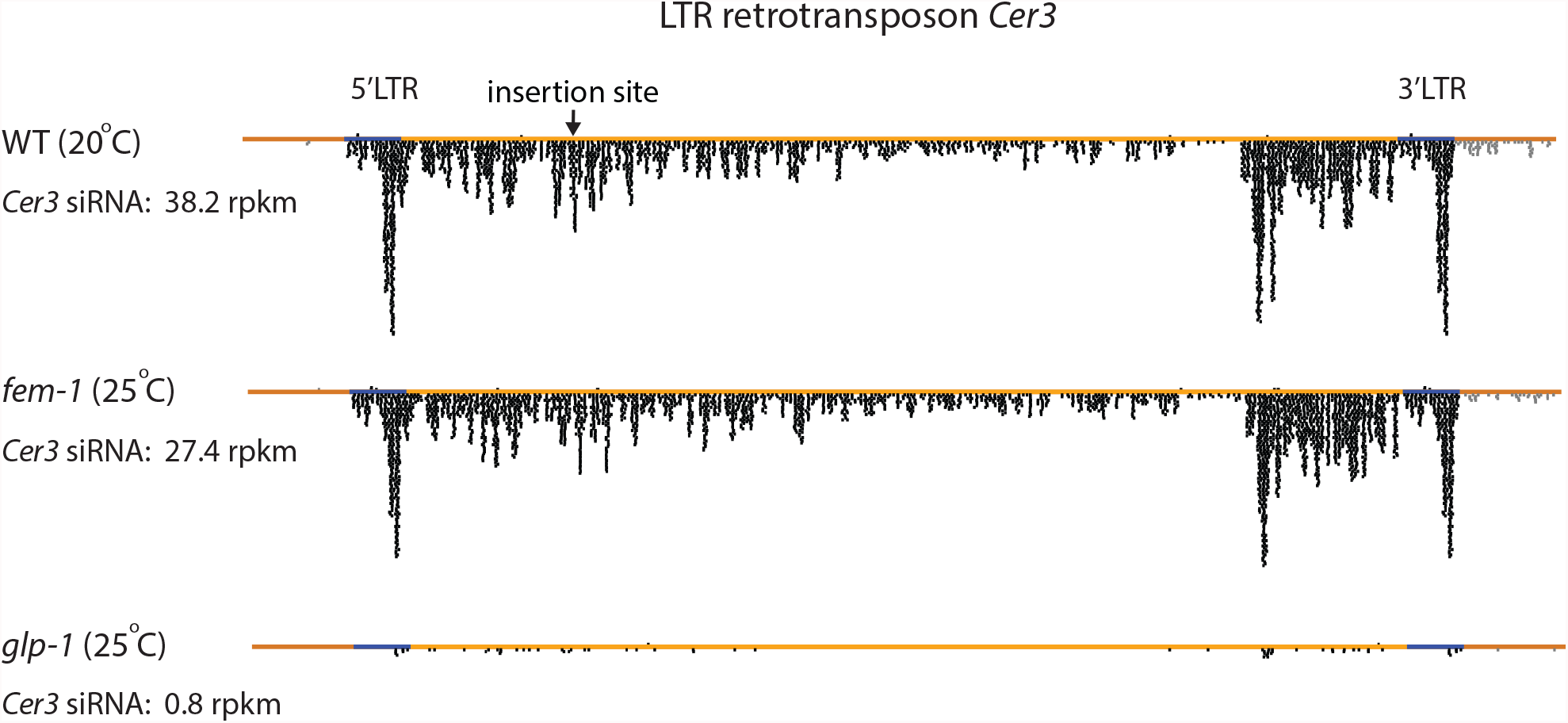
Germline-enriched expression of *Cer3* siRNAs. *Cer3* siRNA track plots for adult WT (20°C), *fem-1*(*hc17*) (25° C, producing functional female germline, but lack of embryo in the uterus due to spermatogenesis defect) [55], and *glp-1*(*e2141)* (25°C, germline depleted) [56] animals. As a quality control for the sRNA-seq, 30%, 12.7%, and 37% of sequenced small RNAs were mapped to microRNAs for WT, *fem-1*, and *glp-1*, respectively. The rpkm values of *Cer3* siRNAs are indicated in the figure. The insertion site in *Cer3* used in this study to express ectopic siRNAs is indicated.

**Fig. S3:**
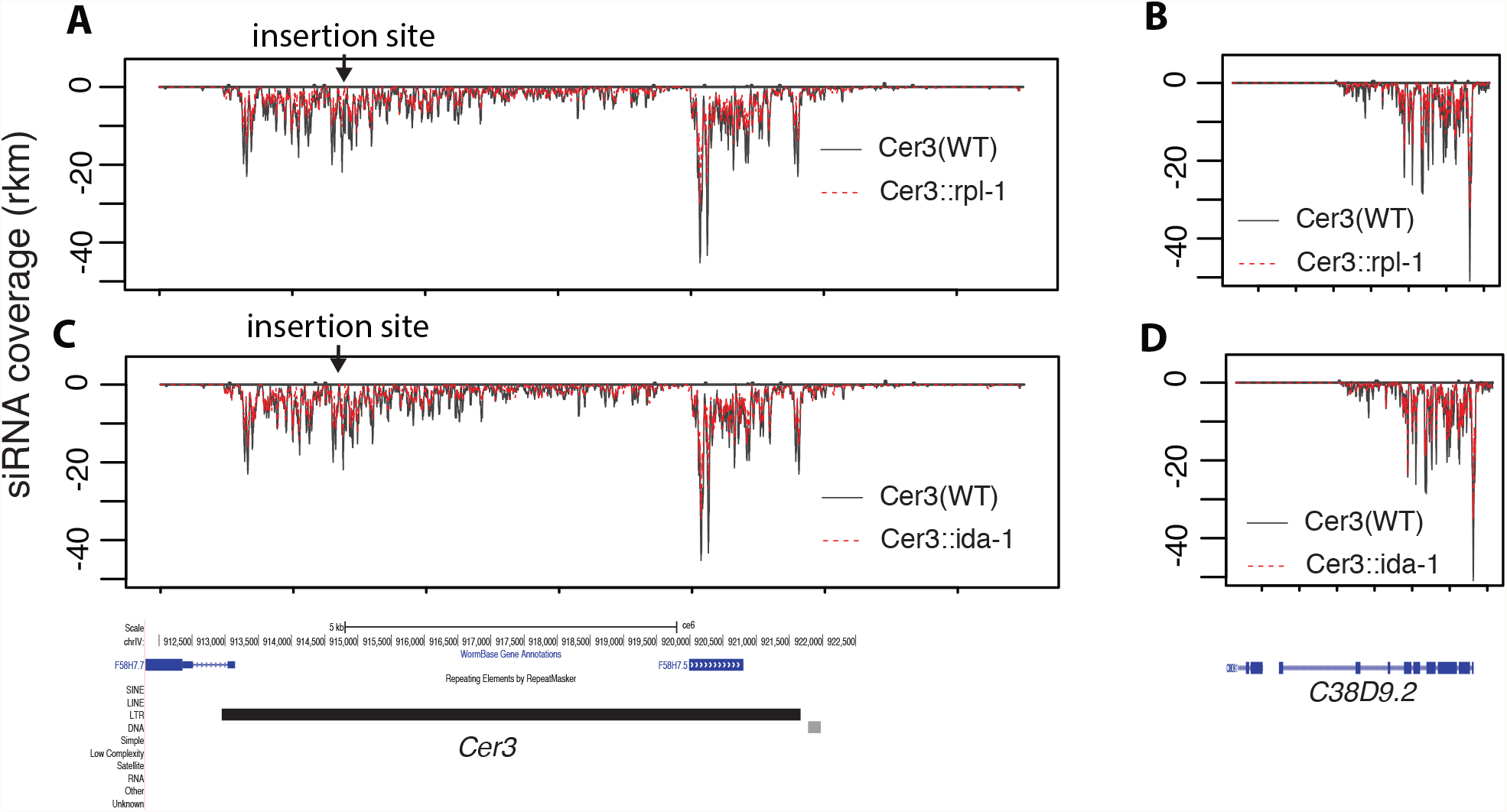
*Cer3* siRNA expression is not affected by insertions. siRNA coverage plots at *Cer3* (**A** and **C**) and *c38d9*.*2* (**B** and **D**) are shown for strains carrying WT *Cer3, Cer3::rpl-1*, and *Cer3::ida-1* as indicated. Sense and antisense siRNA coverages are separately plotted as positive and negative values. WT *Cer3* is used for the alignment and ectopic siRNA coverages are not shown. The WT animals had slightly higher siRNA expressions than the two *Cer3* mutants and other native nuclear RNAi targets, such as *c38d9*.*2* (**B** and **D**). This is likely due to a slight age difference between the samples (data not shown).

**Fig. S4:**
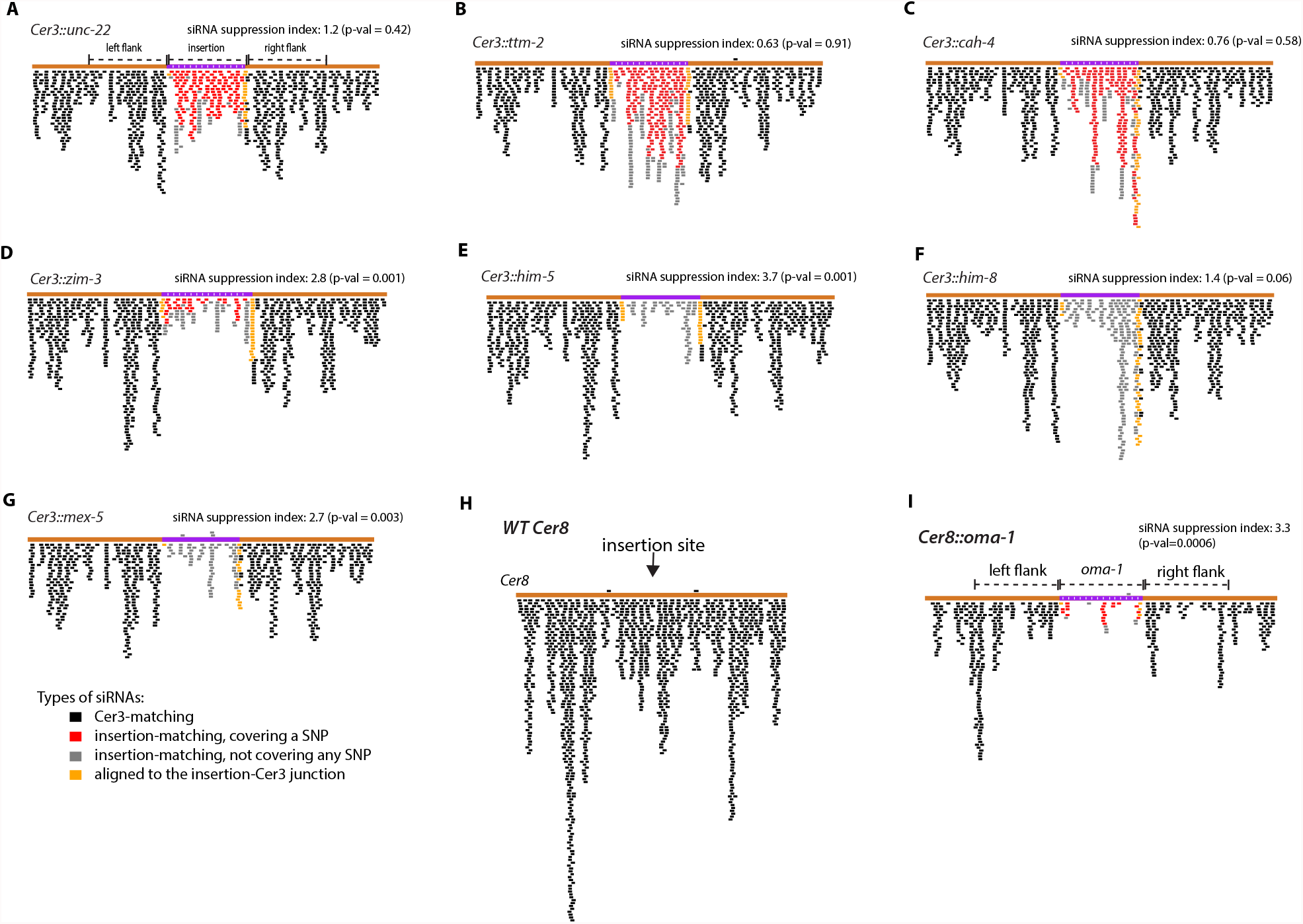
siRNA track plots of additional *Cer3*::insertions (**A**-**G**), WT LTR retrotransposon *Cer8* (**H**), and *Cer8::oma-1* (**I**). Only the 700 nt *Cer8* sequence that flank each side of the insertion site is used for the plots.

**Fig. S5:**
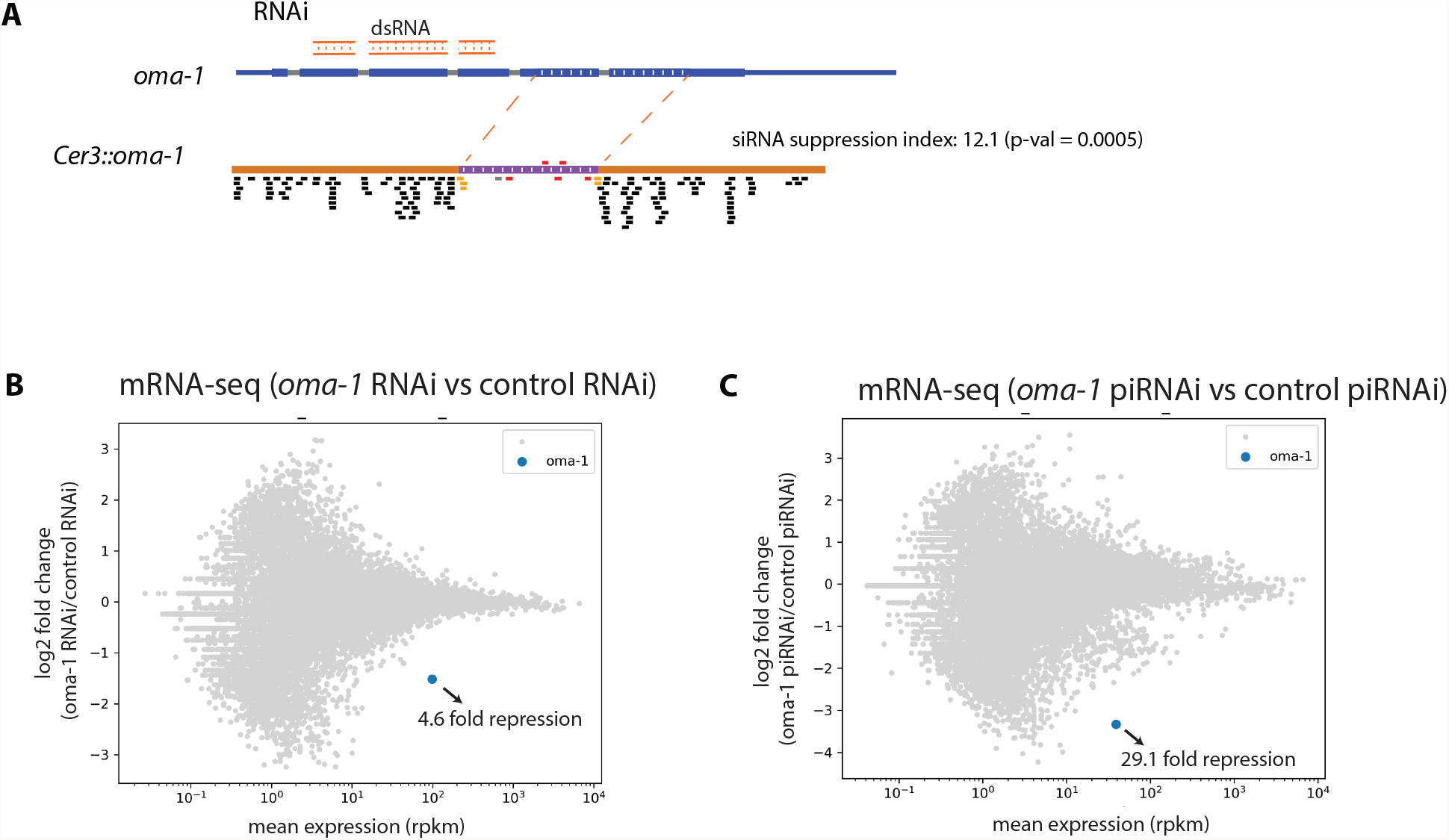
*oma-1* RNAi did not affect siRNA suppression of *Cer3:oma-1*. (**A**): *Cer3::oma-1* siRNA track plot for *oma-1* RNAi or control RNAi (L4440 empty vector) animals. dsRNA targeted region in *oma-1* is indicated. (**B**-**C**): RNA-seq MA-plots of *oma-1* RNAi vs control (L4440) RNAi and *oma-1* piRNAi vs control (*unc-119*) piRNAi, showing that both *oma-1* dsRNA and piRNA led to *oma-1* mRNA repression, but the dsRNA-triggered repression was weaker than the piRNA-triggered repression.

**Fig. S6:**
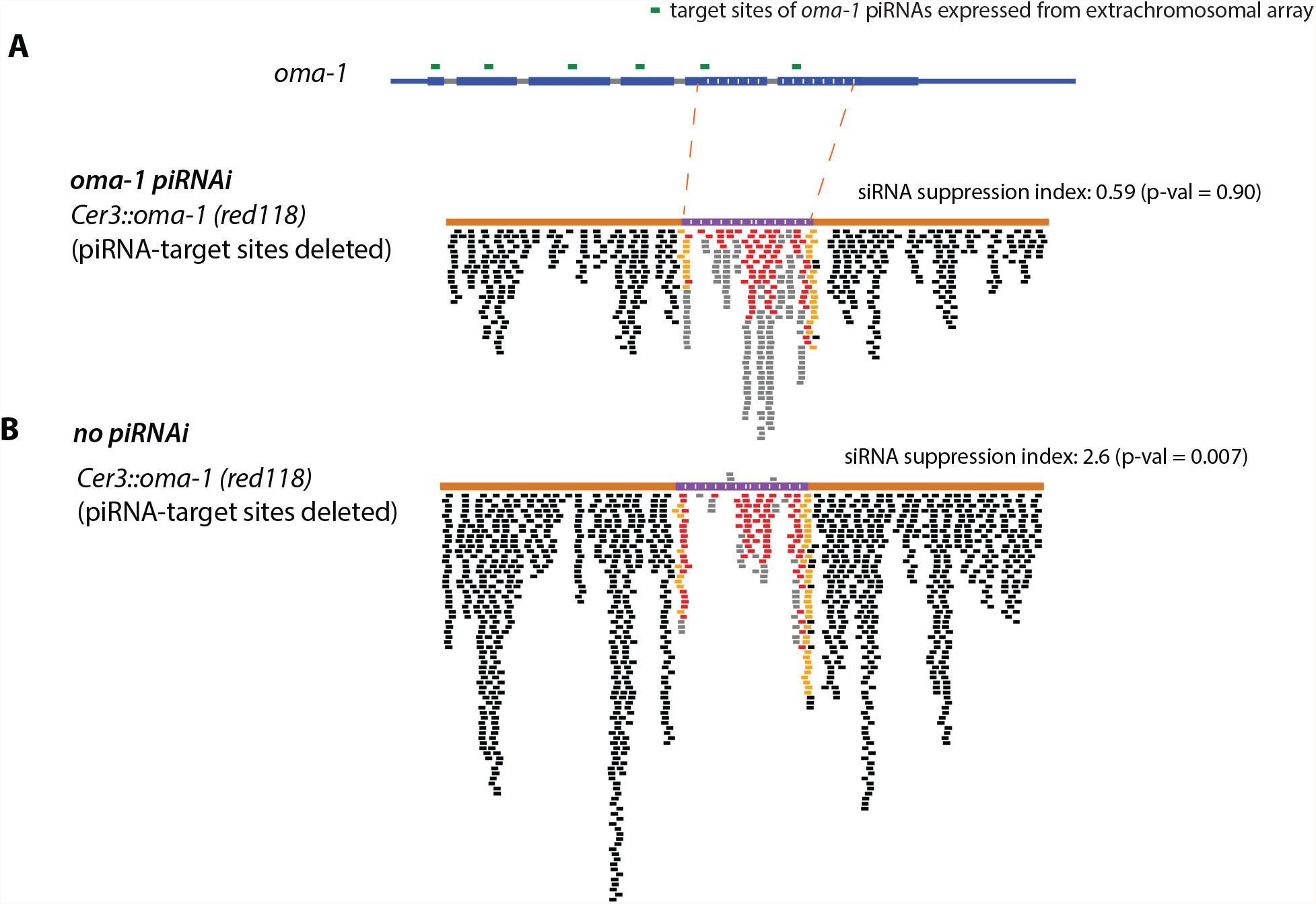
Additional *oma-1* piRNAi experiments. The *oma-1* piRNAi transgene encodes six piRNAs. Their target sites in *oma-1* are indicated in (**A**). Two of the six piRNAs also target the *oma-1* fragment in *Cer3::oma-1* used in this study. To determine whether the effect of *oma-1* piRNAi on siRNA suppression was mediated by the piRNA target sites in the *Cer3::oma-1*, we generated a *Cer3::oma-1* allele (red118) that lacks these two target sites. The siRNA track plots with and without *oma-1* piRNAi are shown in **A** and **B**.

**Fig. S7:**
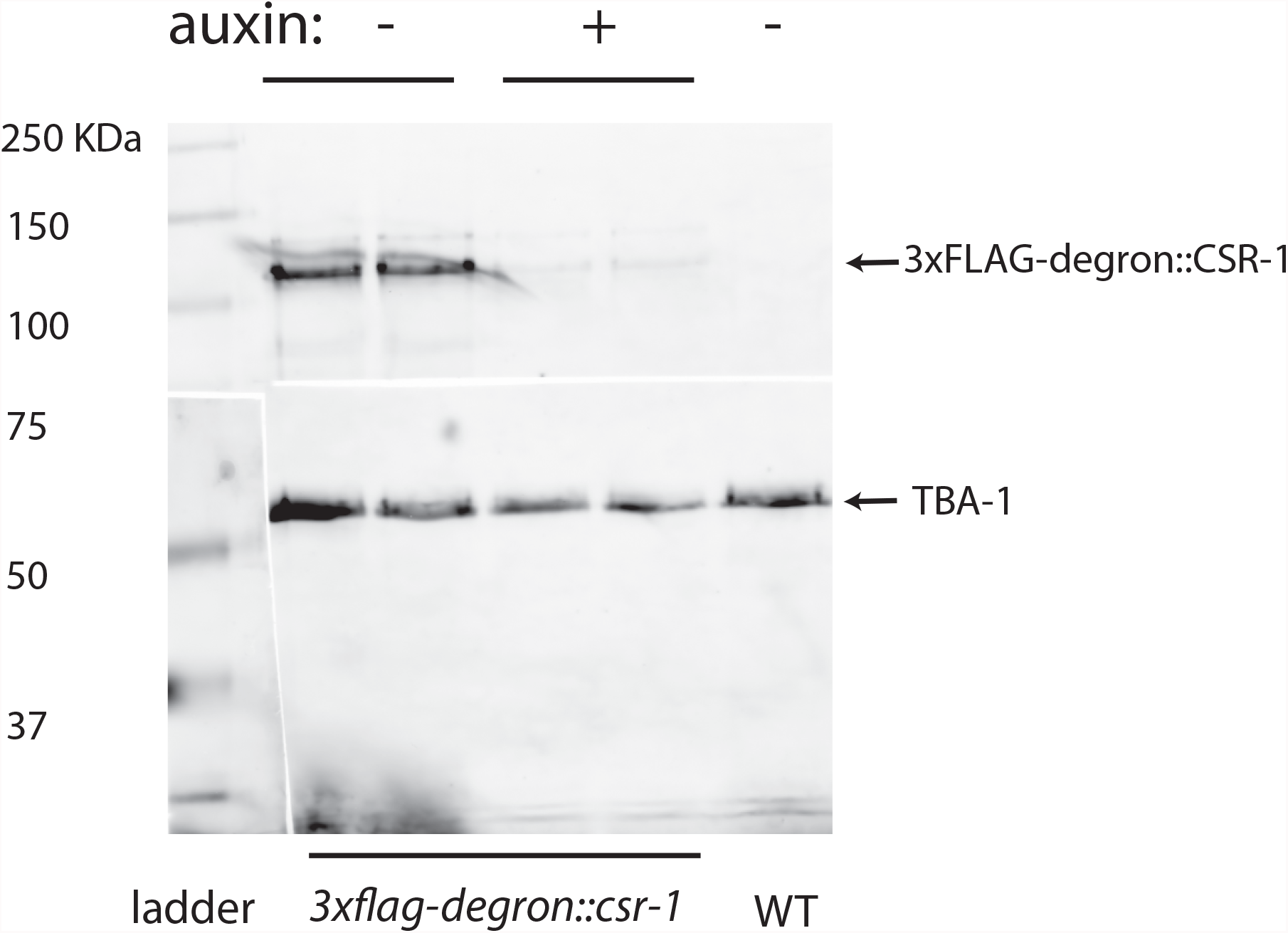
Anti-FLAG western blot of 3xFLAG::degron::CSR-1 confirming auxin-induced degradation (AID) of CSR-1 (87% reduction). We note that, although the CSR-1 depletion was not complete, the animals exhibited a fully penetrant embryonic lethality, a phenotype expected for the loss-of-function *csr-1* mutation (data not shown).

**Fig. S8:**
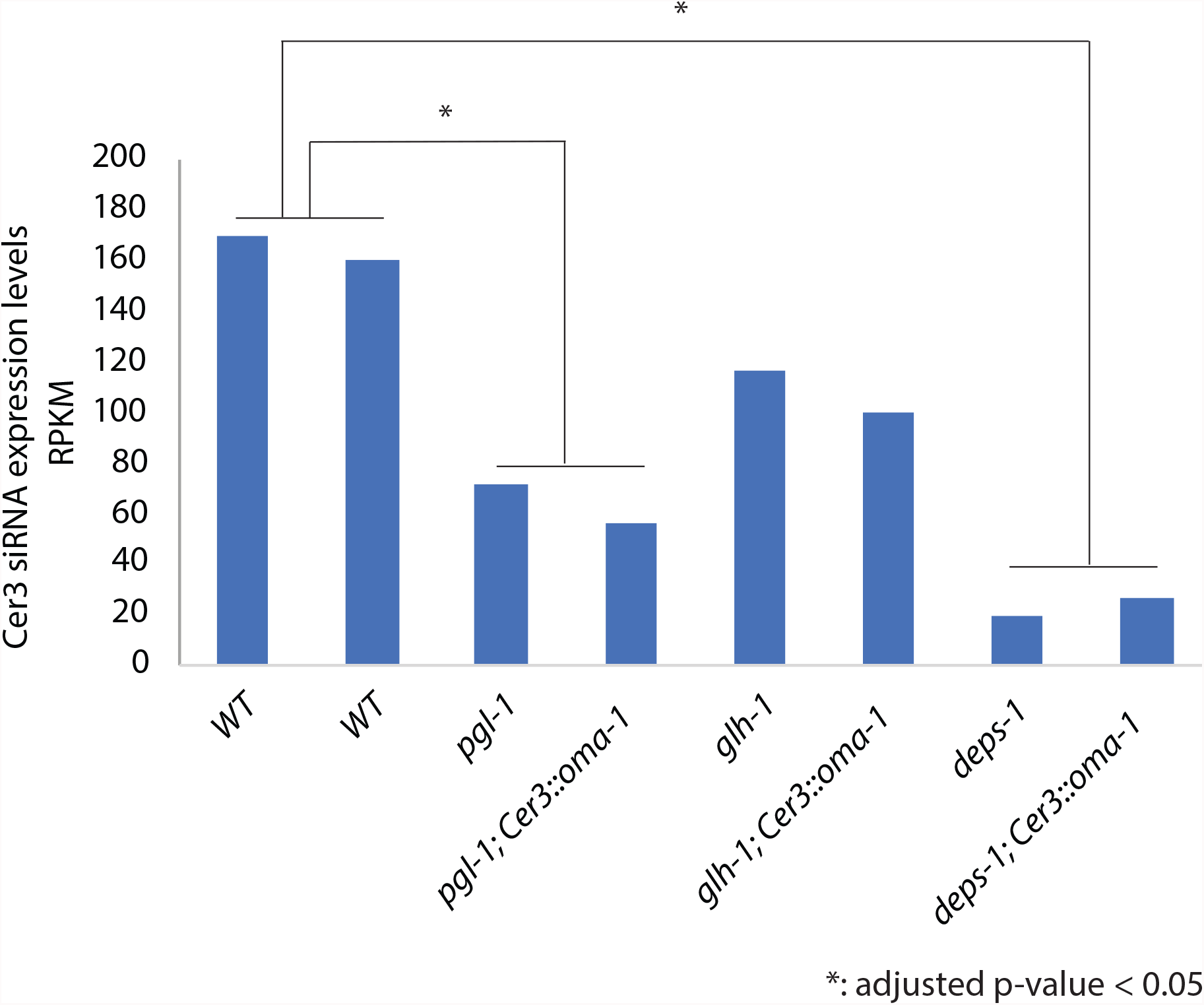
*Cer3* siRNA expression levels in WT and P-granule mutant strains that carry either WT *Cer3* or *Cer3::oma-1*. The full-length WT *Cer3* sequence was used for the alignment to calculate the *Cer3* siRNA levels. DEseq2 [57] was used to calculate the adjusted p-values for the comparison between WT and a mutant background. Both *pgl-1* and *deps-1* mutations were associated with significant reductions in *Cer3* siRNA production (3.7 and 9.2-fold reductions, respectively, adjusted p-values < 1.0×10^−18^). A modest *Cer3* siRNA reduction was observed in the *glh-1* mutant animals (1.7-fold, adjusted p-values = 0.2).

**Fig. S9:**
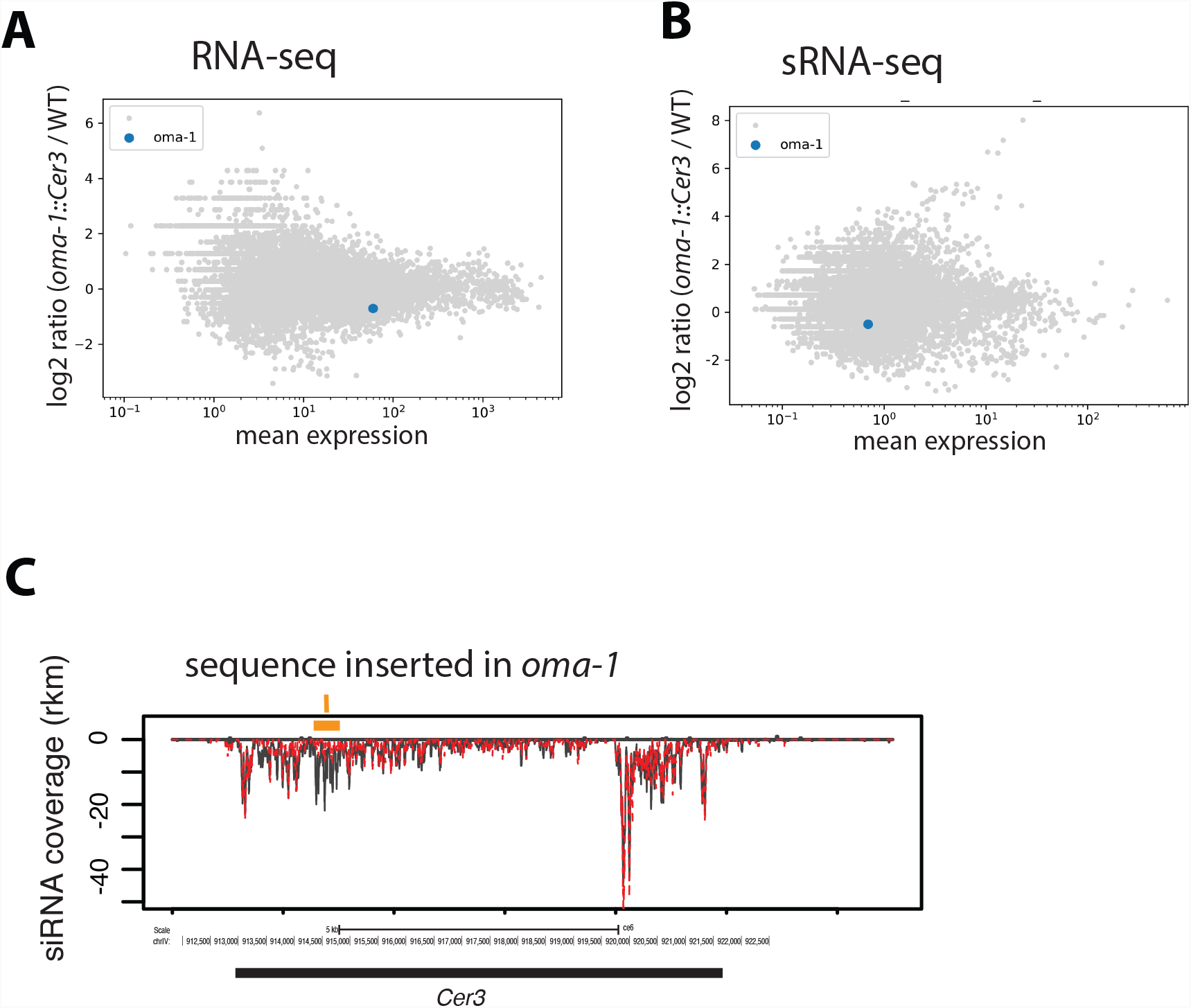
Additional analysis for *oma-1::Cer3* mutant animals. (**A**-**B**) MA-plots comparing *oma-1::Cer3* and WT mRNA (A) and sRNA (B) expressions for all genes, with *oma-1* highlighted in blue. (**C**) siRNA coverage plot at *Cer3* for WT and *oma-1::*Cer3 animals using the same data as Fig. 5A and 5B.

## Methods and Material

### *C. elegans* culture

*C. elegans* were cultured at 20°C on NGM plates seeded with *E. coli* OP50 as described in [58] unless indicated otherwise. Synchronized young adult animals, prepared as described in [40], were used for all experiments in this study.

### CRISPR

CRISPR experiments were conducted using protocols described in [59, 60]. Briefly, the injection mix generally consists of 1 µg/µl Cas9 (IDT), 2.5 µM *dpy-10* sgRNA (Synthego), 0.4 µM ssDNA oligo as *dpy-10* [*cn64*] repair template, 10 µM target sgRNA, target repair template DNA (2 µM for ssDNA oligo or 0.4 µM for dsDNA with single-stranded overhangs generated using a method described in [61]). All CRISPR-generated mutations were confirmed by Sanger sequencing.

### Design of the Cer3-based siRNA generator

An approximately 400 nt cDNA sequence from *C. elegans* protein-coding genes or gfp was inserted into *Cer3* between base positions 914,783 and 914,784 of chromosome IV (WS190). Single-nucleotide mismatches separated by 30 nt intervals were introduced to the inserted sequence to distinguish siRNAs produced from the *Cer3::insertion* locus and ones from native genes.

### Small RNA library preparation and sequencing

Small RNA extraction was performed using the MirVana kit (Thermo Fisher). Small RNA libraries were constructed using the 5’-mono-phosphate-independent, linker ligation-based method as described in [40]. The libraries were pooled and sequenced on the Illumina HiSeq instrument.

### Bioinformatic analysis

All sequence alignments were performed using Bowtie version 1.2.3 [62]. Only the 20-24nt sRNA reads that perfectly aligned to the reference sequence were used for the analysis.

### siRNA track plot, index calculation, and statistics

sRNA-seq reads were aligned to WT or various mutant target genomic region. For siRNA track plots, sequenced sRNAs were drawn based on their genomic alignment locations. Sense and antisense siRNAs were plotted separately above and below the gene track, respectively.

The siRNA suppression index for insertions in *Cer3* was calculated as the ratio between antisense siRNAs density in the 400 nt flanking sequences and the inserted sequence. Note that the ambiguous insertion siRNAs (ones do not cover any SNP position) were included for the calculation. Since some of the ambiguous siRNAs may come from the native gene, the true siRNA suppression index is likely to be higher than the calculated value.

To calculate the statistical significance of the siRNA suppression, we divided the insertion sequence and the 400 bp *Cer3* flanking sequences (left and right) into 50 nt bins. The number of siRNAs matching to each bin were counted. The Wilcoxon Rank Sum Test was performed for these two populations: counts for the insertion bins (G) and counts for the flanking bins (F), with the null hypothesis being G≥F.

### Auxin-induced protein degradation (AID) of CSR-1

The degron-3xflag tag was added to the N-terminus of the longer isoform of CSR-1 (F20D12.1a) by CRISPR in the strain that carries the *Cer3::oma-1(red20)* and *sun-1p::TIR1::mRuby::sun-1 3’UTR* [63]. Synchronized L1 larvae were cultured on plates containing 1 mM auxin or no auxin (as control) until reaching young adults, which were harvested for sRNA-seq. CSR-1 AID was confirmed by both western blotting using the monoclonal anti-FLAG M2 antibody (Sigma) (Fig. S5) and the sterility of auxin-treated animals (data not shown).

### piRNA interference (piRNAi)

*oma-1* piRNAi was induced by an extrachromosomal array carrying the hygromycin-resistance gene and a cluster of piRNA expression units re-coded to target *oma-1* as described in [31]. The control *unc-119* piRNAi transgene was also the same as used in[31]. The extrachromosomal array was selected by hygromycin resistance. The *oma-1* insertion used in the *Cer3::oma-1* (*red20*) allele contains two sites that can be targeted by piRNAs from the transgene. To rule out that the effect on siRNA suppression was due to interaction between *oma-1* piRNAs and *Cer3::oma-1*, we created a new *Cer3::oma-1* allele (*red118*) that deleted the two piRNA-target sites.

### *oma-1* mRNA expression analysis

*oma-1* mRNA levels were measured by either RT-qPCR or RNA-seq as described[64].

## Acknowledgement

We thank Esteban Chen, Helen Ushakov and Elaine Gavin for technical assistance. Research reported in this publication was supported by the Busch Biomedical Grant and the National Institute of General Medical Sciences of the National Institutes of Health under award number R01GM111752 to SG and by KAUST Office of Sponsored Research OSR-CRG2019-4016 to CFJ. Some strains were provided by the CGC, which is funded by NIH Office of Research Infrastructure Programs (P40 OD010440). The content is solely the responsibility of the authors and does not necessarily represent the official views of the National Institutes of Health.

